# Measuring membrane fluidity in live mycobacteria reveals subcellular lateral variation and pole-selective responses to mycomembrane perturbation

**DOI:** 10.1101/2025.09.08.674961

**Authors:** Isabel J. Sakarin, Cyrus Clabeaux, Lia A. Parkin, Jessica C. Seeliger

## Abstract

The cell envelope is an oft-cited factor in the ability of mycobacteria to tolerate antibiotics, host immunity, and environmental stress. *In vitro* studies have led to a prevailing model in which the mycobacterial envelope exhibits low fluidity that hinders the entry of antibiotics and other stressors. Although fluidity affects essential processes and is dynamically regulated across all domains of life, few studies have measured membrane fluidity in live mycobacteria. To address this gap, we used the environmentally-sensitive probe C-Laurdan to develop an imaging- and flow cytometry-based method for measuring cell envelope fluidity directly in live cells. Our approach enables cell envelope labeling across diverse mycobacterial species, including *M. smegmatis* and *M. tuberculosis*. We characterized fluidity as a function of subcellular localization, antibiotic treatment, and genetic perturbation. The unusual growth characteristics of mycobacteria, including polar growth and asymmetric growth and division, contribute to intercellular heterogeneity that is thought to enhance survival under stress. Indeed, we observed that the poles are more fluid than sidewalls, and that the old pole is more fluid than the new pole. Further, daughter cells have unequal membrane fluidity upon division and this asymmetry is reduced in a mutant with decreased asymmetric polar growth. Chemical or genetic disruption of the mycomembrane led to a shared alteration of the fluidity pattern and susceptibility to two antibiotics, suggesting that GP signatures may predict antibiotic susceptibility. This approach expands the toolkit for assessing fluidity in mycobacteria and enables deeper investigation into how biophysical properties influence bacterial physiology and antibiotic susceptibility.

## INTRODUCTION

Lipid membranes define the boundaries of life. Membranes separate cells from their environments and are the sites of essential functions, including nutrient acquisition, energy generation, and environmental sensing. The biophysical properties of membranes are central to their function and are therefore tightly regulated. One such biophysical property is membrane fluidity, a term often used interchangeably with lipid order or packing that we use to refer to the lateral mobility of lipids and proteins within a membrane.

Dysregulation of membrane fluidity interferes with essential cellular functions. For example, membrane fluidity directly correlates with diffusion of protons across the membrane^1,2^ and the binding, insertion, and activity of membrane-associated enzymes^3,4^. Manipulating lipid composition in bacteria to reduce membrane fluidity results in membrane protein mislocalization, impaired cell division, inhibited respiration, dissipation of membrane potential, and activation of cell stress response pathways^5,6^. Fluidity is related to antibiotic sensitivity, as increasing membrane fluidity has been shown to sensitize bacteria to certain antibiotics^7^ and reduced fluidity has been correlated to antibiotic resistance^8^. Fluidity dysregulation is part of the mechanisms of action for some drugs; certain membrane-acting antibiotics specifically disrupt fluid microdomains^9^ or induce the formation of highly fluid domains^10^.

Among bacteria, mycobacteria pose a special case for membrane fluidity. Mycobacteria are distinguished by a cell envelope that is more chemically and structurally complex than that of other bacteria. The mycobacterial envelope includes a unique outer membrane referred to as the mycomembrane. In the most commonly accepted model, this mycomembrane comprises an outer leaflet of diverse free lipids and an inner leaflet of ultra long-chain mycolic acids (MAs; 60-90 carbons) that are covalently linked to underlying layers of arabinogalactan and peptidoglycan. Landmark biophysical studies showed that mycobacterial lipids pack tightly and exhibit extremely low fluidity^11–13^.

Lipid pathways have been implicated in intrinsic resistance to antibiotics. Numerous studies have reported that disruption of genes involved in cell envelope biosynthesis increases sensitivity to some antibiotics^14,15^, and large scale genome-wide chemical genetic interaction studies have identified lipid-related genes as determinants of intrinsic antibiotic resistance in *Mycobacterium tuberculosis (Mtb)*^16–20^. Resistance is frequently ascribed to presumed alterations in the physical properties of the cell envelope, yet this assumption rests on little direct experimental support.

The physical properties of the cell envelope have been studied largely outside of the complex context of live cells. The aforementioned studies demonstrating tight packing were conducted on lipid extracts and cell wall fragments^11–13^. These data have provided the foundation for the current model of mycobacterial membrane fluidity, as well as the overall structure of the mycobacterial cell envelope. More recently, work on protein-free membranes reconstituted into vesicles^21^ and molecular dynamics simulations^22^ suggested that lipid composition determines membrane fluidity and organization. While providing insight into how the unique mycobacterial lipids affect the biophysical properties of membranes, these studies are based on simplified systems and lack full physiological context. For example, reconstituted vesicles lack the covalent linkages to other cell envelope components that are characteristic of mycobacteria. Missing from such analyses are direct measurements of membrane properties in live bacteria for which the composition and structure of the cell envelope remain fully intact.

To address this gap, we turned to membrane-targeting fluorophores to assess envelope properties without disrupting their native state. In particular, we focused on environmentally-sensitive fluorophores, for which emission properties change in response to the local lipid environment. While diverse environmentally-sensitive, membrane-targeting fluorophores are available to measure membrane properties such as polarity, fluidity, and domain formation in live cells^23–26^, these have not been broadly applied to mycobacteria. Membrane-targeting fluorophores were screened for their distribution in cells^27^ and metabolically-incorporated polarity-sensitive dyes were developed as diagnostic reagents^28,29^, but these studies did not explore the potential to report on membrane properties. We chose to focus on Laurdan-based probes because they (1) have been characterized extensively in model and cellular membranes; (2) provide generalized polarization (GP) as a quantitative metric commonly interpreted as membrane fluidity, and (3) only require non-polarized emission measurements (vs. lifetime or anisotropy), making the method broadly accessible^30^. Laurdan GP has been reported in mycobacterial studies, but only in vesicles reconstituted from mycobacterial lipids^21^ or in cells in bulk^31^.

Given the precedent for using Laurdan probes in live cell, we posited that they could bridge the gap between mycobacterial cell biology and membrane biophysics. We developed an imaging- and flow cytometry-based method for measuring cell envelope fluidity in live mycobacteria using the carboxylic Laurdan variant C-Laurdan. Our approach robustly labeled the cell envelope of diverse mycobacterial species, including *M. smegmatis* and the human pathogens *M. abscessus* and *M. tuberculosis*. By characterizing patterns of fluidity, we found that cell envelope fluidity varies within and between cells. Mycobacteria grow asymmetrically to yield daughter cells with different sizes and growth rates, and the resulting heterogeneity is thought to promote survival, e.g., in response to antibiotics^32,33^. Consistent with this asymmetry, we observed that cell envelope fluidity is not equal in daughter cells upon division, and this heterogeneity is reduced in a mutant with decreased asymmetric polar growth (Δ*lamA*). We found that membrane fluidity differs spatially between cell poles and sidewalls, and this pattern is altered after chemical or genetic perturbation of the mycomembrane. Further results combining chemical treatment and genetic disruption suggested GP signatures as potential predictors of antibiotic susceptibility. Overall, our method is a broadly applicable tool for probing mycobacterial membrane fluidity in live cells and opens new avenues for understanding how envelope properties relate to bacterial physiology and antibiotic susceptibility.

## RESULTS

### C-Laurdan labels Mycobacterium smegmatis

We developed a microscopy-based method to measure membrane fluidity in live mycobacteria. We tested three Laurdan-based probes in *M. smegmatis* (*Msm*): (1) Laurdan, (2) the carboxylated variant C-Laurdan^34,35^, and (3) Pro12A, a derivative designed for reduced internalization in mammalian cells^36^ (Figure 1A). Each dye was incubated with live *Msm* in growth medium, then washed and imaged. We found that incubating cells with C-Laurdan for 5 min was sufficient to yield robust, reproducible, and dose-dependent fluorescence, as detected by confocal microscopy (Figure 1B). C-Laurdan signal localized to the cell periphery, with punctate staining adjacent to this periphery and also within the cell body. In contrast, labeling with Laurdan or Pro12A was comparatively weak and diffuse (Figure 1B). These initial observations supported C-Laurdan as a candidate for labeling mycobacterial cell envelope membranes.

**Figure 1.**
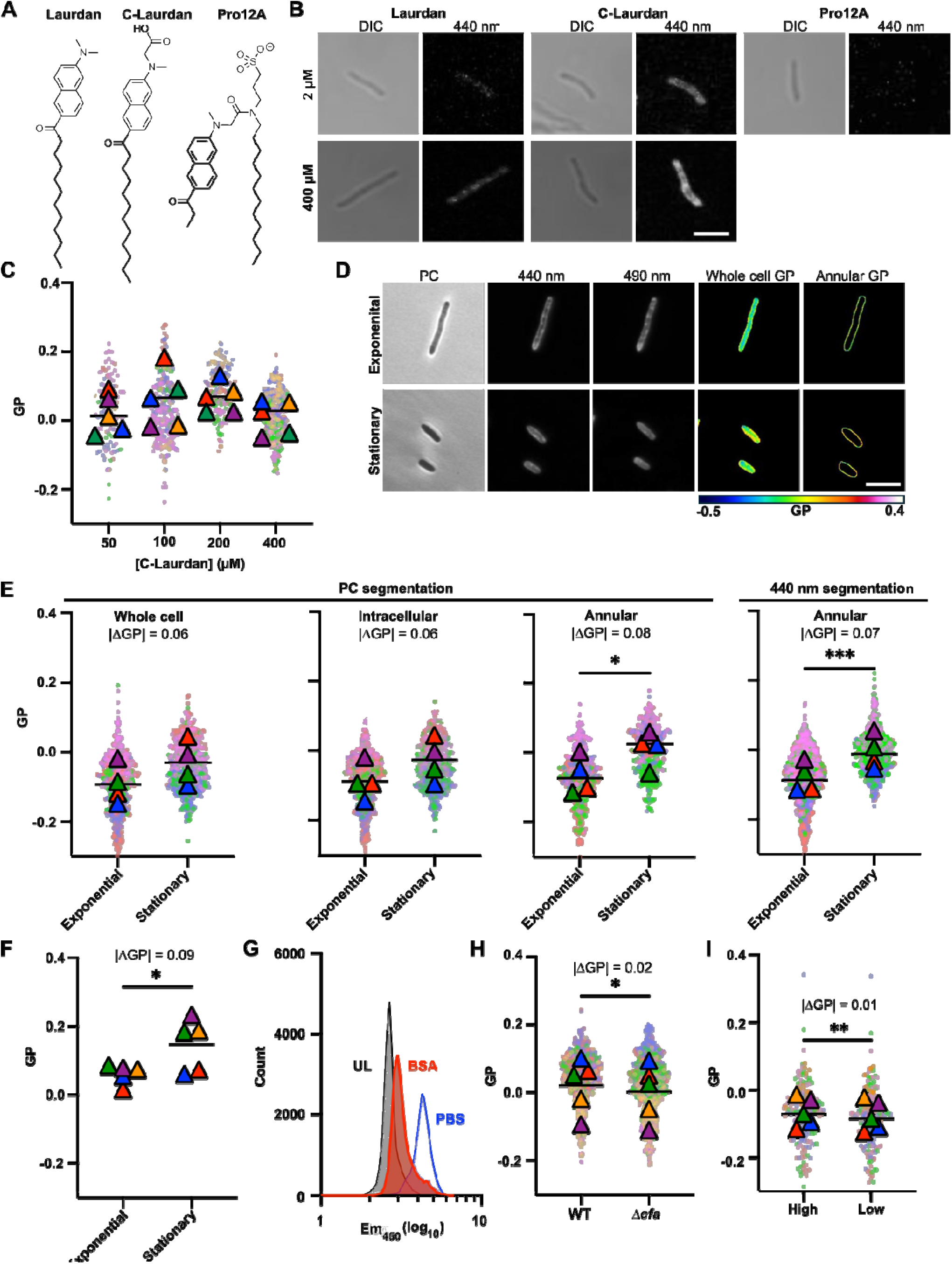
C-Laurdan reports on changes in membrane properties. **A.** Structures of Laurdan, C-Laurdan, and Pro12A. **B.** Representative confocal microscopy images of wild-type (WT; strain mc^2^155) *M. smegmatis* (*Msm*) labeled with Laurdan, C-Laurdan, or Pro12A. Differential interference contrast (DIC) and 440 nm emission images are shown. Scale bar = 5 µm. **C.** Average annular GP of WT *Msm* after 5-min labeling with 50, 100, 200, or 400 µM C-Laurdan. The means (triangles) are superimposed on individual data points (circles) from N[=[5 independent experiments (colored by replicate; ∼50 cells per replicate) and were used to calculate *p*-values, means (black line), and ΔGP. **D.** Representative confocal images of WT *Msm* from exponential or stationary phase cultures. Phase contrast (PC), fluorescence emission, and generalized polarization (GP) maps of the entire cell (whole cell) and signal from the cell periphery (annular). **E.** Average GP of exponential or stationary phase WT *Msm* cells analyzed for whole cell, intracellular, or annular GP segmented based on the phase contrast image, and annular GP segmented based on 440 nm emission. N = 4, ∼100 cells per replicate. **F.** Average GP of exponential or stationary phase WT *Msm* cells measured by flow cytometry. N = 5, each with data from 20,000 events. **G.** Flow cytometry of unlabeled (UL, grey) *Msm and* C-Laurdan-labeled cells treated with either BSA (red) or PBS (blue). **H.** Average annular GP of WT or Δ*cfa Msm.* N = 5, ∼100 cells per replicate. **G.** Average annular GP of the high-GP or the low-GP side of individual *Msm* cells when bisected along the long axis. N = 5, ∼20-50 cells per replicate. **H.** Data in C were analyzed by ordinary one-way ANOVA. Data in E, F, H, and I were analyzed using paired two-tailed *t-*tests; \**p* ≤ 0.05, \*\**p* ≤ 0.01*, ***p* ≤ 0.001. Unless otherwise noted, cells were labeled with 400 µM C-Laurdan for 5 min and segmentation was based on the phase contrast.

We next assessed whether C-Laurdan perturbs cell envelope fluidity when it integrates into membranes, especially at 400 µM, which is higher than concentrations reported for other fluorescent membrane stains in mycobacteria^27,37,38^. We found that annular GP was independent of C-Laurdan concentration (Figure 1C). Across all concentrations, we observed that a subset of cells was unlabeled, but that this subset was smallest when labeling with 400 µM C-Laurdan. Therefore, for all subsequent experiments, we used 400 µM C-Laurdan and excluded unlabeled cells from GP analysis.

### C-Laurdan generalized polarization detects expected changes in membrane properties

The emission maximum of Laurdan probes such as C-Laurdan responds to environmental factors. The properties and uses of Laurdan are detailed in a recent review by Gunther et al.^30^ Briefly, in more polar, disordered, and hydrated environments, emission is red-shifted. Laurdan’s environment-sensitive emission profile is used to define the metric generalized polarization (GP) as (I_₄₄₀_□–□I_₄₉₀_)/(I_₄₄₀_□+□I_₄₉₀_), where I_₄₄₀_ and I_₄₉₀_ are fluorescence intensity at wavelengths that represent the response of C-Laurdan to lipid-ordered and lipid-disordered environments, respectively. GP is a commonly used ratiometric proxy for membrane fluidity.

To validate that our method reports on changes in membrane fluidity, we tested whether C-Laurdan GP reflected shifts expected from conditions in which cell envelope changes are well established. Growth phase-dependent transitions provide a relevant model, as bacteria undergo characteristic metabolic and morphological shifts upon entry into stationary phase^39,40^. This transition is also associated with alterations in lipid composition^41–43^ and increased antibiotic tolerance^43–46^. We therefore compared membrane fluidity between exponential and stationary phase cells.

We first established that in the growth medium used (Middlebrook 7H9 supplemented with casamino acids; see Methods), *Msm* grows exponentially up to an optical density at 600 nm (OD_600_) of ∼3 before plateauing at OD_600_ ∼4. Cells from exponential (OD_600_ 1-2.5) and stationary phase (OD_600_ 4-6) cultures exhibited distinct cell sizes, with the latter distinctly shorter, as has been previously noted^47^ (Figure 1D). Although these conditions give rise to distinct morphologies, C-Laurdan labeling yielded similar patterns of staining that were consistent with the peripheral and punctate staining observed in initial experiments with exponential-phase cells (Figure 1D).

The annular pattern suggested that C-Laurdan labels the membranes of the cell envelope while the additional punctate signal in the interior supported membrane-proximal and cytoplasmic labeling of other structures. In principle, these nanometer-scale differences cannot be discriminated by visible wavelength light microscopy. However, given these visible qualitative distinctions within individual cells, we asked whether different image segmentation approaches would affect the absolute and relative GP values of exponential and stationary phase cells. We analyzed images by quantifying whole cell GP, isolating intracellular GP, and isolating annular GP after segmentation based on the phase contrast image (Figure 1E). Across all three analyses, stationary phase cells showed higher average GP values than exponential phase cells, indicating greater membrane rigidity. However, |ΔGP| was larger (0.08 vs. 0.06) and reached statistical significance (*p* < 0.05) only in the annular GP analysis, suggesting that isolating the annular signal preserves distinct information pertinent to the cell envelope. To increase accessibility of our method for labs without phase-contrast imaging capabilities, we also tested segmenting cells based on 440 nm emission. This approach yielded comparable results (Figure 1E, |ΔGP| = 0.07, p ≤ 0.001). Given this similarity and for technical reasons, we used phase contrast and 400-nm segmentation interchangeably throughout the rest of the study.

We also developed a flow cytometry assay as a complementary approach that enables analysis of whole-cell GP far more rapidly than by microscopy (∼20,000 cells/min by flow vs. ∼200 cells/hour by microscopy). Exponential and stationary phase cells exhibited similar |ΔGP| values when measured by either method, indicating that microscopy provides an unbiased and representative sampling of the population despite its lower throughput (Figure 1F). While absolute GP values are not expected to match across methods due to differences in optical configurations (e.g., filters and detectors), the agreement in trend supports the validity of both approaches. We favor microscopy for its ability to provide spatial information, which we leveraged in subsequent analyses. Overall, the magnitude of the difference between stationary and exponential phase cells is similar to those reported as biologically meaningful in bulk assays on mycobacteria and other bacteria^31,48,49^. Overall, this comparison of supports C-Laurdan detection of physiologically significant differences in membrane fluidity.

Our microscopy results support C-Laurdan labeling of membranes in the cell envelope. Since mycobacteria have two membranes, C-Laurdan may localize to the plasma membrane, mycomembrane, or both. To assay for mycomembrane localization, we tested whether C-Laurdan was extractable and therefore surface-accessible by incubating C-Laurdan-labeled *Msm* with bovine serum albumin (BSA) and measuring fluorescence emission by flow cytometry (Figure 1G). Incubation with BSA resulted in decreased fluorescence compared to PBS-treated controls, but higher fluorescence than unlabeled cells (Figure 1E). This may reflect incomplete removal of C-Laurdan from the mycomembrane under the conditions used, or indicate that C-Laurdan is also localized to the plasma membrane. While we cannot rule out the latter, this result nonetheless indicates that C-Laurdan incorporates into the mycomembrane.

In an effort to determine whether C-Laurdan GP reports on plasma membrane fluidity, we compared annular GP between the Δ*cfa* mutant and its corresponding WT parent strain. Loss of *cfa* function results in the accumulation of phospholipids with unsaturated fatty acid tails instead of the methylated tuberculostearic acid^50^. This perturbation is expected to alter lipid packing and thereby membrane fluidity specifically in the plasma membrane. Indeed, Δ*cfa* has a lower annular GP than the WT parent strain (Figure 1H). However, the |ΔGP| of 0.02, while statistically significant, is smaller than differences we observed in the growth phase experiments, and the biological relevance of GP changes on this scale remains unclear. Other systems that introduce well-characterized changes to specifically the plasma membrane are needed to determine more conclusively if C-Laurdan localizes to this membrane as well as the mycomembrane. As a negative control for biological significance, we compared the average annular GP of the high-GP and the low-GP side of individual *Msm* cells when bisected along the long axis, a comparison in which there is no literature precedent for a biological difference (Figure 1I). The magnitude of the difference was small (|ΔGP| = 0.01) but statistically significant (*p* ≤ 0.01). This suggests that small GP differences, even when statistically distinguishable, may not reflect meaningful biological changes. Given the unresolved ambiguity regarding plasma membrane labeling, we interpret GP values in this study as reporting on overall cell envelope membrane fluidity.

### C-Laurdan is broadly applicable to pathogenic *Mycobacteria* and *Nocardia* species

Based on the rapid and robust labeling of *Msm* by C-Laurdan, we then asked whether this approach could be applied to a broader range of pathogenic actinobacteria. We found that no further optimization of our method was necessary to label *M. tuberculosis* (*Mtb*) mc^2^6020 Δ*panCD* Δ*lysA*, *M. abscessus* (*Mab*) 390R, *Mab* 390S, *M. chelonae, M. kansasii, M. fortuitum, M. avium, M. scrofulaceum,* and *N. farcinica* (Figure 2A). For all species tested the pattern of labeling was similar, with peripheral and intracellular punctate signal (Figure 2A). Although the number of replicates does not permit statistically rigorous comparison, the cell envelope of *Mtb* is qualitatively the most rigid (Figure 2B). While the more important takeaway is the generality of the method, an initial observation for follow up is that fast-growing species trend towards having more fluid cell envelope membranes than slow-growing species.

**Figure 2.**
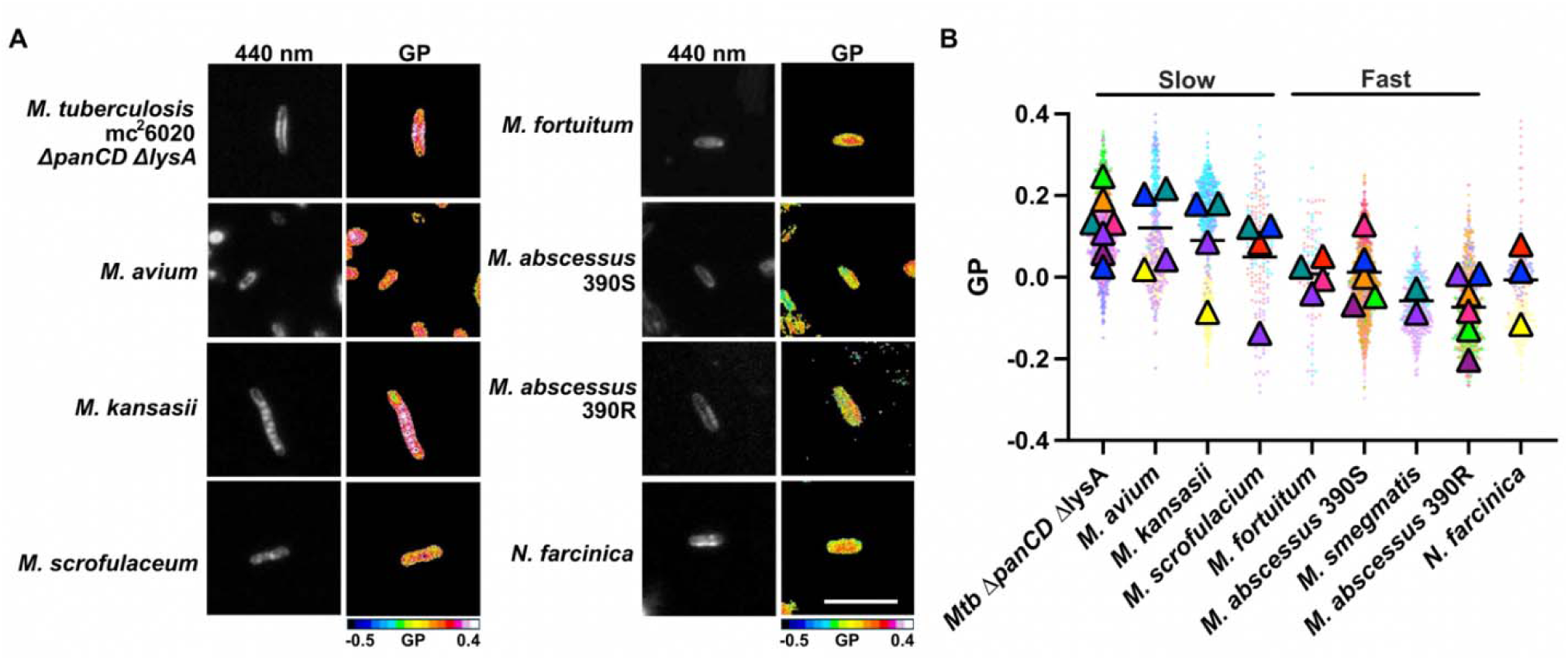
C-Laurdan labeling is generally applicable to *Mycobacteria* and *Nocardia* species that cause human infections. **A.** Representative confocal images and GP maps of *Mycobacteria* and *Nocardia* species labeled with 400 µM C-Laurdan for 5 min. Scale bar = 5 µm. **B.** Annular GP of slowing-growing mycobacteria (*M. tuberculosis (Mtb)* mc^2^6020 Δ*panCD* Δ*lysA, M. avium* subspecies *avium* Chester, *M. kansasii, M. scrofulaceum*), fast-growing mycobacteria (*M. fortuitum, M. abscessus* 390R, *M. smegmatis, M. abscessus* 390S), and *Nocardia farcinica.* Data are shaded by independent experiment. Segmentation for GP analysis was based on 440 nm emission.

### Membrane fluidity varies both between and within individual cells

GP maps revealed visually distinct patterns of membrane fluidity both within and between cells (Figure 3A). During replicative growth, mycobacteria elongate at the poles^47,51^, which are sites of active cell envelope synthesis^52^. Also, the inner membrane domain, a physically and biochemically distinct plasma membrane fraction enriched in cell wall biosynthetic enzymes, localizes to these polar regions^38,53,54^. Given this lateral organization of envelope biosynthesis, we hypothesized that membrane fluidity would likewise exhibit spatial variation along the cell axis. Indeed, we found that poles are more fluid than sidewalls (|ΔGP| = 0.03, Figure 3B), suggesting that newly synthesized membranes differ in fluidity from older membrane regions.

**Figure 3.**
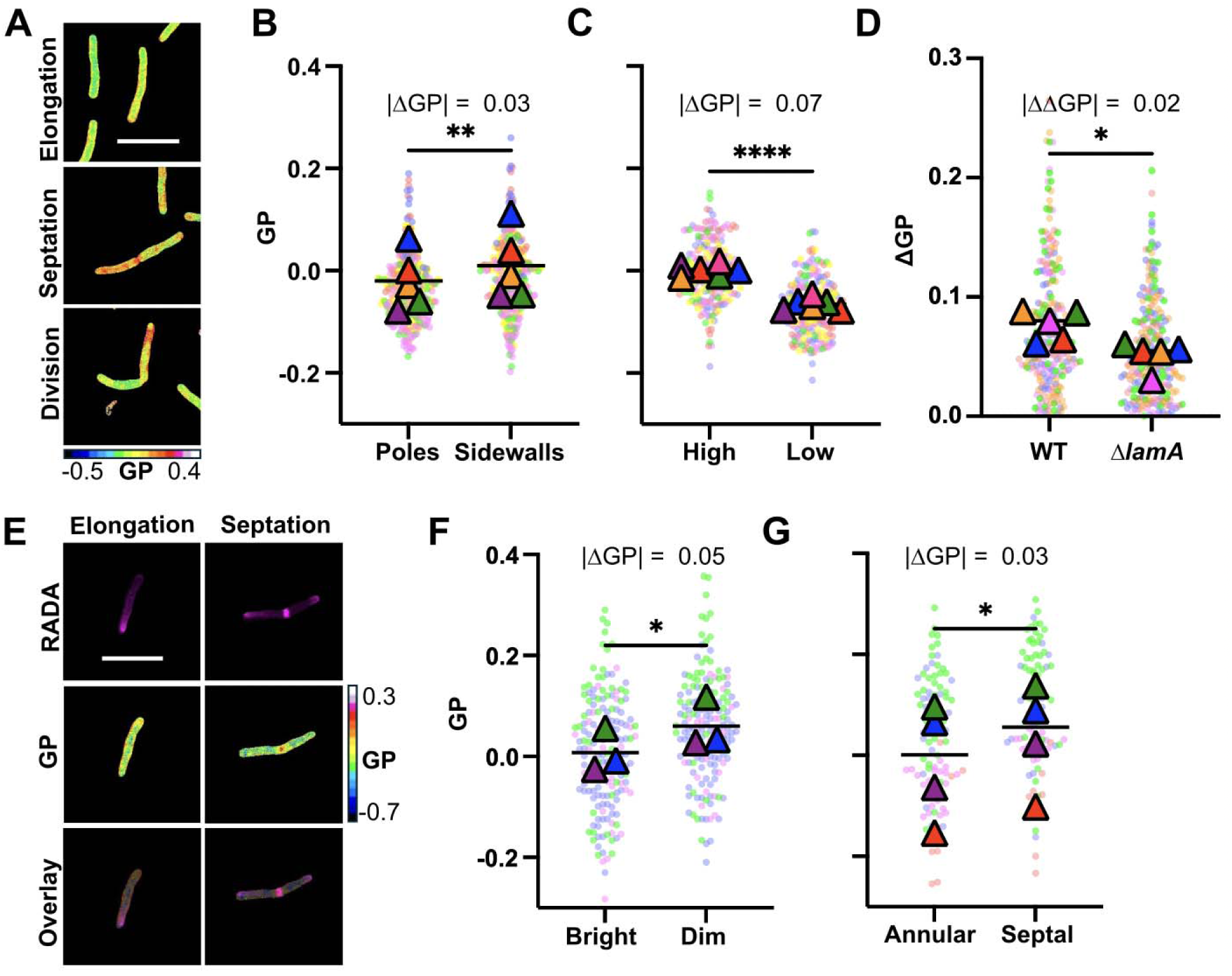
Membrane fluidity varies both between and within individual cells. **A.** Representative GP maps of WT *Msm* highlighting cells in different growth stages based on the presence of a septum or observation of v-snapping (division). Scale bar = 5 µm. **B.** Average annular GP measured at the poles or sidewalls of WT *Msm*. Data from N□=□5 independent experiments, ∼100 cells per replicate. **C.** Average annular GP of high-GP vs. low-GP daughter cells in pairs of dividing cells from WT *Msm*. N = 6, ∼100 cells per replicate. **D.** The average difference in annular GP (ΔGP) between paired high-GP vs. low-GP daughter cells of WT or Δ*lamA Msm*. N = 5, ∼100 cells per replicate. **E.** *Msm* was metabolically labeled with 25 µM RADA for 15 min and then labeled with 400 µM C-Laurdan for 5 min. Representative RADA images and GP maps are shown. **F.** Average annular GP at the RADA-bright or RADA-dim pole. N = 3, ∼50 cells per replicate. **G.** Annular vs. septal GP based on identification of the septum by RADA labeling. N = 4, ∼20 cells per replicate. Statistical analysis: B, C, F, G: Paired two-tailed *t-*test. D: Paired one-tailed *t-*test. * *p* ≤ 0.05, ** *p* ≤ 0.01, \*\*\**p* ≤ 0.001. For all panels, segmentation for GP analysis was based on phase contrast.

Mycobacteria also grow and divide asymmetrically, yielding daughter cells with different sizes and growth rates^32^. The resulting intercellular heterogeneity is thought to promote survival, as the resulting daughter cells demonstrate differential resistance to stressors, including the first-line anti-tuberculosis drugs rifampicin and isoniazid^32,33^. Given the dominant model linking membrane properties to cell envelope function and antibiotic resistance, we asked whether this physiological and functional asymmetry is reflected in fluidity. We compared the average annular GP between daughter cells in dividing cell pairs (identified by v-snapping) and found that cell envelope fluidity differs significantly between daughter cells upon division (|ΔGP| = 0.07, Figure 3C). We further hypothesized that this daughter cell heterogeneity would be reduced in the mutant Δ*lamA*, which exhibits decreased asymmetric polar growth and altered antibiotic susceptibility^33^. Indeed, the difference in GP between daughter cells was significantly reduced in Δ*lamA* (|ΔΔGP| = 0.02, Figure 3D, ΔGP calculated as GP_high GP daughter_ – GP_low GP daughter_), supporting a model in which *lamA*-dependent growth asymmetry contributes not only to size heterogeneity, but also to membrane fluidity heterogeneity, with possible functional consequences for stress tolerance.

In light of our evidence suggesting that envelope biosynthesis locally alters membrane fluidity (Figure 3B), we hypothesized that the old pole, where active cell wall synthesis is concentrated^32,55^, would exhibit distinct membrane fluidity compared to the new pole. To test this, we used the fluorescent D-amino acid analog rhodamine D-alanine (RADA) as a fiducial marker of peptidoglycan synthesis, allowing us to distinguish the old (RADA-bright) pole from the new (RADA-dim) pole (Figure 3E). We found that in co-labeled cells, the GP is significantly lower at the RADA-bright pole than the RADA-dim pole, indicating that polar growth rate correlates with increased fluidity (Figure 3F). RADA also marks the septa, which have significantly higher GP and are thus less fluid than the surrounding cell envelope (Figure 3G). These results suggest that elongation and septation, though both involving active cell envelope synthesis, give rise to distinct membrane states.

### Chemical treatment affects membrane fluidity and results in localized fluidity changes

While the molecular targets of many antibiotics are well characterized, little is known about how these treatments affect the broader physiology and organization of bacterial cells. Understanding how antibiotics impact cell envelope properties, such as membrane fluidity, may provide insights into their distinct mechanisms of action and the resulting bacteriostatic and bactericidal effects. We first aimed to characterize the effect of ethambutol, a cell wall-acting antibiotic. Ethambutol inhibits polymerization of arabinan domains in arabinogalactan, thereby reducing attachment sites for mycolic acids in the mycomembrane. We hypothesized that this biochemical disruption would lead to measurable changes in cell envelope fluidity. Treatment with ethambutol at approximately 1x and 20x MIC^56–58^ for 3 h resulted in increased average annular GP and thus reduced fluidity compared to vehicle-treated controls (Figure 4A, 4B). Further, the increase in GP was most pronounced at the poles, whereas the change at the sidewalls was not significant (Figure 4B). This localized effect was also reflected in demographs mapping fluidity along the long axis as a function of cell size, as a proxy for progress in the cell cycle, and in average line profiles (Figure 4C, 4D). From these analyses we also noted that 3 h ethambutol treatment resulted in cell shortening (Figure 4C).

**Figure 4.**
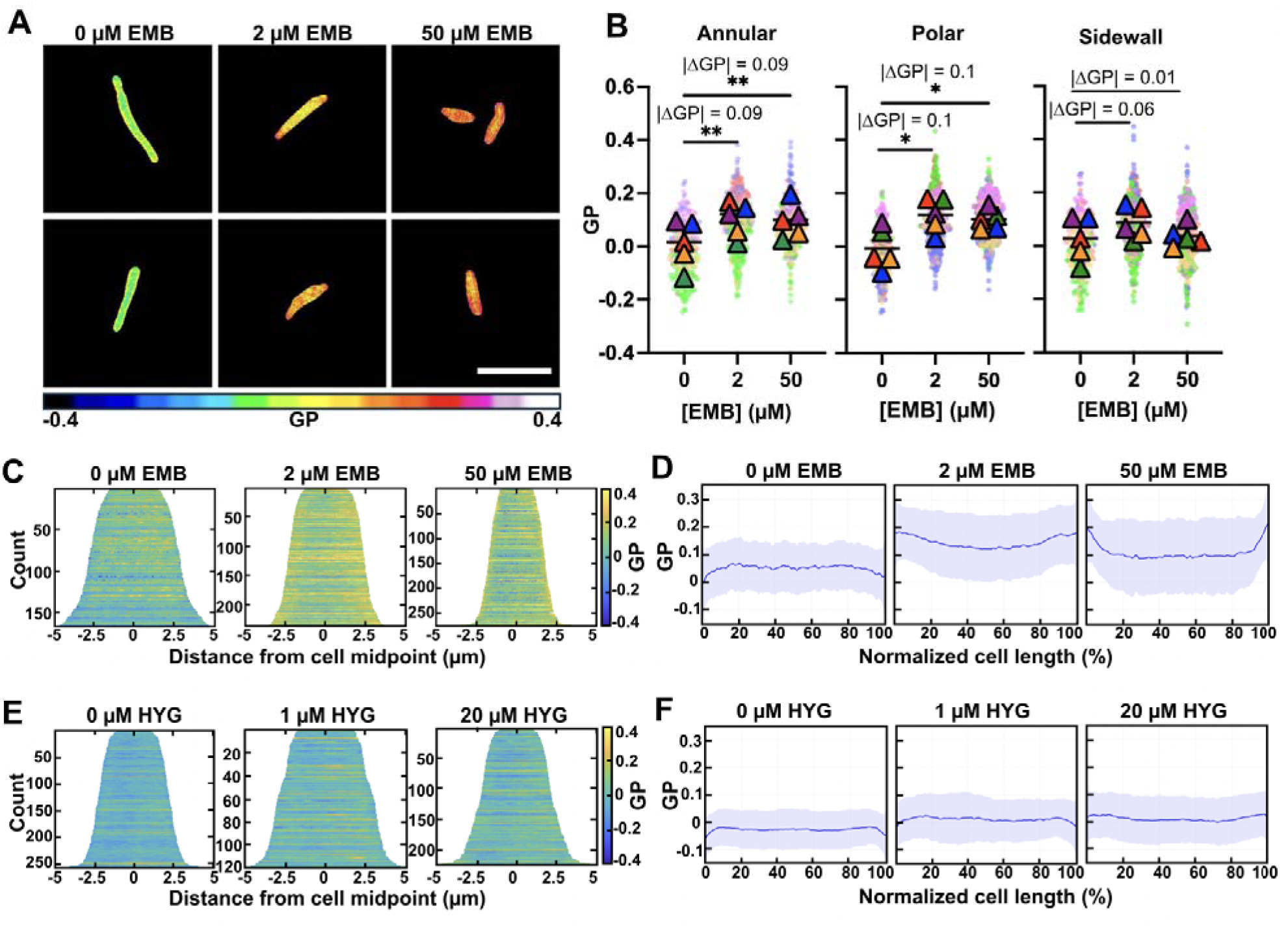
Chemical treatment affects membrane fluidity and results in localized fluidity changes. **A.** Representative GP maps of WT *Msm* treated with 0, 2, or 50 µM ethambutol (EMB) for 3 h prior to C-Laurdan labeling. Scale bar = 5 µm. **B.** Average, polar, and sidewall annular GP of vehicle and EMB-treated WT *Msm*. N = 5 independent experiments, ∼100 cells per replicate. **C, D, E, F:** Whole cell GP line profiles were extracted and used to create (C, E) demographs and (D, F) average line profiles after treatment with EMB (C, D: N = 4) or hygromycin (E, F: N = 3) for 3 h. Statistical analysis: Repeated measures ANOVA with Tukey’s multiple comparisons. * *p* ≤ 0.05, ** *p* ≤ 0.01. For all panels, segmentation for GP analysis was based on phase contrast signal.

We then asked whether this decrease in polar fluidity was specific to ethambutol as a cell wall synthesis inhibitor or characteristic of a general response to antibiotic treatment. As a comparator we used the protein synthesis inhibitor hygromycin. After analogous treatment with approximately 1x and 20x MIC for 3 h, the GP did not change markedly at the poles or across the entire cell axis (Figure 4E, 4F). This suggests that the localized fluidity changes caused by ethambutol treatment are related to the inhibition of cell wall assembly, rather than from a nonspecific stress response to antibiotic exposure at and above minimum inhibitory concentrations.

### LprG-rv1410c loss of function and ethambutol treatment yield similar patterns of GP and antibiotic susceptibility

Given that fluidity has been hypothesized to play a role in cell envelope function and intrinsic antibiotic resistance, we sought to assess membrane fluidity in a mutant with defective cell envelope function. The LprG-Rv1410c pathway is highly conserved across all mycobacteria, required for virulence^59,60^ and associated with triacylglyceride (TAG) transport^60^. Disruption of the *lprG-rv1410c* operon increases susceptibility to multiple antibiotics and small molecules^16,18,61–63^ and increases the accumulation of fluorescent dyes^63^.

These studies have interpreted these changes as a result of disruption of the cell envelope upon loss of LprG-Rv1410c function. We sought to test this assumption by measuring cell envelope membrane fluidity. The annular GP of Δ*lpxrG-rv1410c* is significantly higher than for WT and therefore the cell envelope is more rigid (|ΔGP| = 0.06, Figure 5A). We examined GP along the long axis and found that Δ*lprG-rv1410c* cells have increased GP at the poles compared to sidewalls (Figure 5B). This polar rigidity is similar to what we observed after ethambutol treatment (Figure 4D).

**Figure 5.**
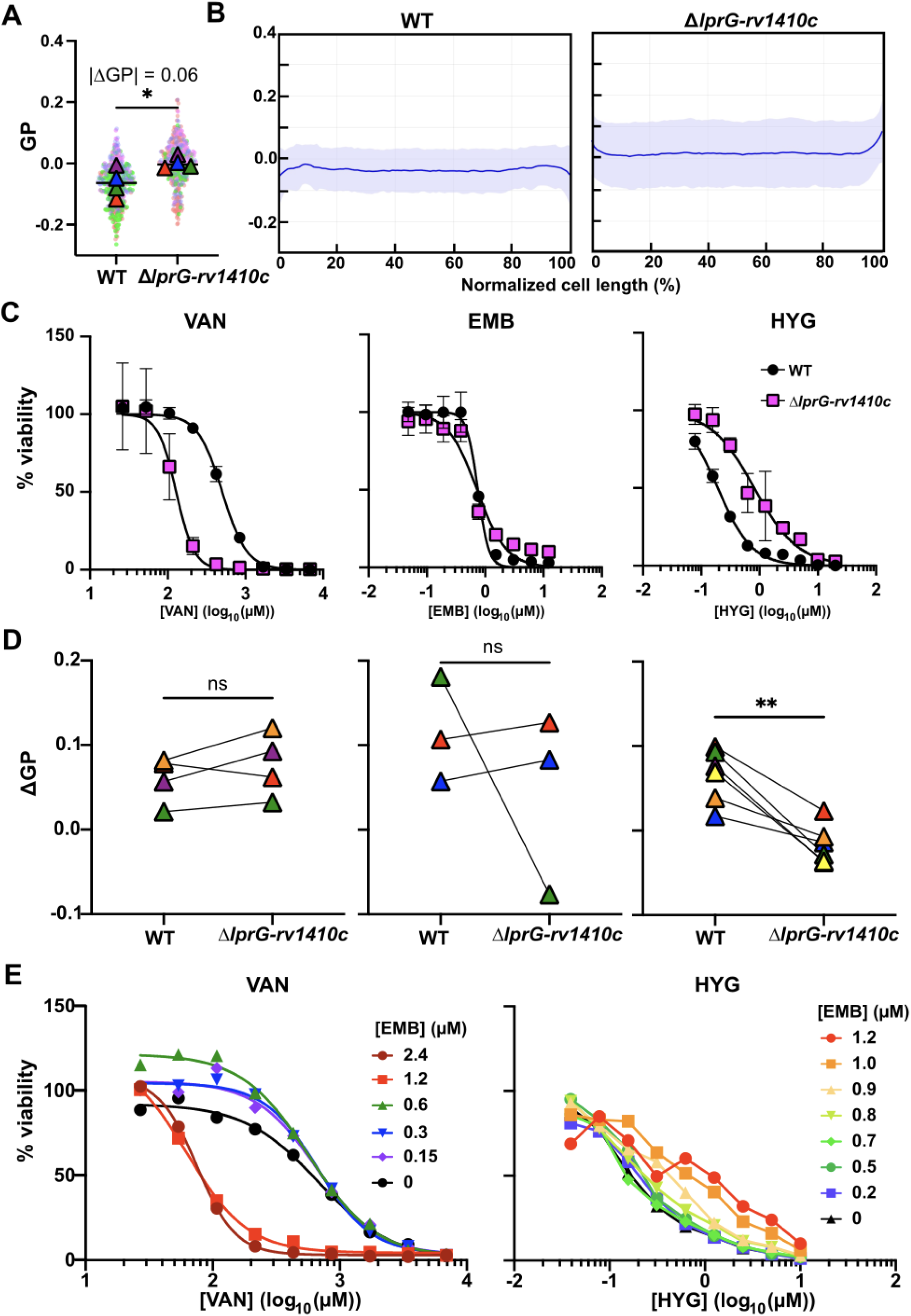
LprG-rv1410c loss of function and ethambutol treatment yield similar patterns of GP and antibiotic susceptibility. **A.** Average annular GP of WT and Δ*lprG-rv1410c Msm.* N = 4, ∼100 cells per replicate. Statistical analysis: paired two-tailed t-test, \**p* ≤ 0.05. **B.** Whole cell GP line profiles were extracted and used to create average line profiles. A, B: Segmentation for GP analysis was based on the phase contrast signal. **C.** Dose response curves for autoluminescent WT (black circles) and Δ*lprG-rv1410c* (pink squares) *Msm* treated with vancomycin (VAN), EMB, or hygromycin (HYG). **D.** ΔGP (calculated as average annular GP_0 abx_ – GP_+abx_) for WT and Δ*lprG-rv1410c Msm* after treatment with 20 µM HYG B, 50 µM EMB, or 5 mM VAN for 3 h. Data are colored by independent replicates with lines connecting paired replicates. Segmentation based on the 440 nm emission. **E.** Dose response curves for autoluminescent WT *Msm* after co-treatment with EMBand either VAN or HYG. For VAN, data averaged from N = 2 experiments were fit using a four-parameter variable-slope model (log[inhibitor] vs. response). For HYG, substantial variability in the data precluded fitting to a nonlinear curve, so data averaged from N = 4 experiments are connected to guide the eye.

We evaluated the antibiotic dose response of the Δ*lrpG-rv1410c* strain and observed that, relative to wild type, the mutant displayed heightened resistance to hygromycin, similar susceptibility to ethambutol, and increased susceptibility to vancomycin (Figure 5C). We asked how GP changes after treatment with these antibiotics. We treated WT and Δ*lprG-rv1410c* with 20 µM hygromycin, 50 µM ethambutol, or 5 mM vancomycin (∼20x MIC) for 3 h and compared the change in GP (ΔGP), calculated as the difference between average annular GP without antibiotic and with antibiotic treatment (Figure 5D). The WT and mutant strains displayed a range of responses to antibiotics as measured by GP, both in the direction and magnitude of the change. The observation most clearly correlated with changes in antibiotic susceptibility was the relative magnitude of the change in GP. The mutant was more susceptible to vancomycin and responded with a greater change in membrane fluidity than the WT. In contrast, the mutant was less perturbed by hygromycin, to which it is more resistant. This result is a tantalizing suggestion that relative antibiotic susceptibility is reflected in the degree of GP perturbation, but further studies will be needed to elucidate this.

We also noted that both chemical and genetic disruptions expected to perturb the mycomembrane yielded similar patterns of membrane fluidity, with increased rigidity at the poles. We hypothesized that this might indicate a shared response in membrane properties that would correlate with the antibiotic susceptibility profile under these conditions. Therefore, we tested whether ethambutol treatment of WT *Msm* would recapitulate the response of Δ*lprG-rv1410c* to vancomycin and hygromycin. Indeed, ethambutol treatment increased resistance to hygromycin in a dose-dependent manner (Figure 5E). Inversely, ethambutol increased susceptibility to vancomycin in a dose-dependent manner. While preliminary, these results provide a tantalizing sugestion that GP signatures have the potential to predict antibiotic susceptibility.

## DISCUSSION

We developed an imaging-based method for measuring cell envelope fluidity with the environmentally-sensitive probe C-Laurdan. We found that while Laurdan and Pro12A labeled *Msm* dimly and diffusely, the carboxylic variant C-Laurdan labeled along the cell periphery, indicating envelope membrane staining (Fig. 1B). Labeling was achieved without permeabilization, an unexpected advantage for obtaining physiological information compared to the established requirement for permeabilization prior to labeling Gram-negative bacteria with Laurdan and other membrane probes^6^.

Explicit charges are expected to influence the association and internalization of membrane probes by increasing the energetic barrier to crossing lipid bilayers. In eukaryotic cells, this is illustrated by the limited internalization of C-Laurdan, which exists partly in an anionic state at physiological pH, and the outer leaflet specificity of Pro12A, which has a fixed negative charge^36^. Mycobacteria are reported to have negative surface charge^64^, which may explain why the fully anionic Pro12A fails to label mycobacteria, and why a probe bearing a partially anionic population can label more strongly than the uncharged Laurdan. The development of organelle-targeted Laurdan derivatives highlights the potential to design probes that selectively label membranes based on biological function^65^. Overall, the greater chemical diversity afforded by the growing array of modified Laurdan variants offers an opportunity to empirically assess mycobacterial membrane labeling, with the potential for improved characteristics such as higher signal-to-noise and specificity for the cell envelope.

In addition to labeling the cell periphery, C-Laurdan also stained punctate regions adjacent to the cell periphery and within the cell body (Fig. 1B, 1D). Puncta of intracellular lipid labeling are often referred to as intrabacterial lipid inclusions (ILIs), although this term is most commonly used to refer to larger and more central structures induced by specific conditions such as low nitrogen and nutrient limitation^66^. While we did not analyze these structures and their associated GP in this study, intracellular staining by C-Laurdan is a direction for future investigation, as ILIs have been implicated in bacterial metabolism, stress response, and virulence^66^.

As expected, we found that the cell envelope fluidity differs in exponential versus stationary phase cells, possibly reflecting growth phase-associated changes in lipid composition^41–43^. We found that stationary phase *Msm* cells have less fluid membranes, indicating that the changes accompanying stationary phase promote tighter lipid packing, reduced membrane hydration, and/or restricted movement of membrane lipids and proteins (Figure 1C, 1D). Interpreting this decrease in membrane fluidity in the context of lipidomic findings from *M. tuberculosis*, we note that stationary phase cells showed increased abundance of TAGs and a shift toward higher-mass TAG species^41^. While these are observations made in *Mtb,* we would expect such changes to be shared with *Msm,* which also has these lipids classes. TAGs are found in the mycomembrane^67^, and our fluidity data is consistent with a model in which these neutral, hydrophobic lipids localize between leaflets to increase packing of the mycomembrane. Interestingly, in *Mtb,* stationary phase cells exhibit an increase in saturated fatty acyl content in all major glycerophospholipid subclasses – phosphatidylethanolamine, phosphatidylinositol, and cardiolipin – that are found in the plasma membrane^41^. Increased fatty acyl saturation is predicted to increase fluidity and decrease packing^68,69^ – the opposite of what we observed by C-Laurdan GP. This provides additional, albeit circumstantial, evidence that C-Laurdan reports predominantly on the mycomembrane.

Our finding that BSA extracts C-Laurdan fluorescence is another, more direct indication that C-Laurdan labels the mycomembrane (Fig. 1E). Using the Δ*cfa* mutant as a tool, we were unable to conclude whether C-Laurdan in the plasma membrane contributes significantly to the observed GP, as the |ΔGP| of 0.02 between WT and Δ*cfa* is smaller than other differences reported in this study and is comparable in magnitude to the difference observed between the two sides of a cell when bisected along its long axis – a scenario for which there is no literature precedent for a biological difference. If this |ΔGP| is not biologically meaningful, this result indicates either that loss of *cfa* has minimal impact on membrane fluidity, or that C-Laurdan GP primarily reflects the mycomembrane rather than the plasma membrane. In light of this remaining ambiguity, we are working to develop alternative approaches to selectively measure mycomembrane fluidity.

In addition to *Msm*, our approach robustly labeled several actinobacteria that cause human infections. The slow-growing species^70^ – *Mtb, M. avium, M. kansasii, and M. scrofulaceum* – qualitatively have more rigid membranes than the fast-growing species^70^ – *M. abseccus, M. fortuitum*, *M. chelonae,* and *Msm.* This trend in membrane fluidity among fast- and slow-growers correlates with the different classes of mycolic acids found in the two groups. The slow growers have the longest mycolic acids among actinobacteria, with chain lengths in the range of 70-90 carbons^71^. Increasing chain length has been shown to correlate with decreased diffusion of glycolipids within the mycomembrane, implying lower fluidity^72^. Additionally, the presence of highly modified and oxygenated mycolates (keto-, methoxy-, and wax mycolates) in these species may contribute to mycomembrane rigidity^71^. The fast growers contain α’ mycolates as a major component of their mycolic acid profiles and these are the shortest among mycobacterial mycolates (∼60 carbons) and may therefore contribute to the comparatively higher fluidity of these species^71^.In contrast to the diverse mycomembrane lipid compositions across actinobacteria, the core architecture of the plasma membrane is thought to be well conserved across species^73–75^. The fact that we detect GP differences across species is further circumstantial evidence that the method reports on mycomembrane fluidity.

We found that cell envelope fluidity is not equal in daughter cells upon division, and this heterogeneity is reduced in a mutant with decreased asymmetric polar growth (Δ*lamA*). Further analysis of spatial differences – such as examining the GP at the septa of dividing cells and the variation in GP along the long axis of high-vs. low-GP daughter cells – could uncover trends related to membrane remodeling, asymmetric division, and inheritance of distinct membrane properties.

With respect to our observation that envelope membrane fluidity is distinct at the poles vs. sidewalls, other studies have speculated on this intracellular variation, albeit with contrasting conclusions. On the one hand, polar increases in mycolyltransferase activity (inferred from turnover of a substrate analogue) led Wuo et al. to propose that the poles are more fluid than the sidewalls, although the reason for connecting enzyme activity to membrane fluidity is not clear^76^. On the other hand, the apparent sequestration of lipoglycans at the poles by Smelyansky et al. suggested that the poles are less fluid, thus preventing newly synthesized lipoglycans from diffusing across the cell^77^. Our results support the former idea that the poles are more fluid and also that the old pole is more fluid than the new pole. C-Laurdan labeling could be combined with metabolic and chemical labeling methods to determine more directly the correlation between membrane fluidity and enzyme activity and lipid localization, as reported by those tools. Such information could help define differences in composition that underlie differences in fluidity.

We note that the poles and septa, while both regions of active cell envelope synthesis, are more fluid and more rigid, respectively, compared to the overall cell envelope. A study that simultaneously tracked peptidoglycan and mycomembrane synthesis with metabolic labels revealed that these processes are coordinated distinctly at the poles vs. septa^78^. The higher membrane rigidity at the septum may serve to resist stretch and deformation during cell division, as peptidoglycan is fractured and then repaired^78^.

We found that treatment with ethambutol at approximately 1x and 20x the MIC for 3 h resulted in increased annular GP, indicating increased rigidity (Figure 4A, 4B). This contrasts with a previous report showing that similar conditions of dose-dependent ethambutol treatment promoted the movement of a metabolically incorporated fluorescent trehalose glycolipid^72^.

Several factors could explain this apparent discrepancy. First, diffusion was measured by fluorescence recovery after photobleaching (FRAP), which has been shown to cause membrane damage that could alter fluidity in the area of analysis^79^. Second, FRAP was directed to the sidewall, whereas our approach measured the GP of the entire cell as well as poles and sidewalls. Sidewall GP indicated lower fluidity after ethambutol treatment, but this difference was smaller than at the poles and not statistically significant, so these results do not necessarily contradict the FRAP study.

Our results challenge the expectation that inhibiting mycomembrane biosynthesis will increase membrane fluidity, with an inferred increase in permeability. Also, observed fluidity changes are not uniform, but affect poles more than sidewalls. We found that these effects are unique to ethambutol compared to the translational inhibitor hygromycin, but shared with loss of function in the lipid transport pathway genes *lprG-rv1410c*. We found a further correlation with antibiotic susceptibility, as ethambutol treatment and *lprG-rv1410c* loss of function induced similar changes in susceptibility to vancomycin and hygromycin. These initial observations suggest correlations between mycomembrane perturbation, subcellular patterns of membrane fluidity, and the profile of antibiotic susceptibility. Further studies will be necessary to test the generality of the observed distinctions and whether they correlate with inhibitor mode of action and gene function.

The reduced fluidity of Δ*lprG-rv1410c* complements a previous report of increased surface rigidity in this mutant measured by atomic force microscopy^63^. While stiffness and fluidity probe distinct aspects of the envelope, AFM stiffness has been correlated with membrane microviscosity in other systems^80^.The concordance of these measurements in Δ*lprG-rv1410c* suggests a potential coupling between membrane fluidity and surface rigidity that merits further study.

Overall, C-Laurdan labeling is a broadly applicable tool for measuring membrane fluidity in live mycobacteria that opens new avenues for understanding how cell envelope dynamics contribute to physiology and phenotypic properties. The characterization presented here lays the groundwork for using fluidity measurements to explore mycobacterial envelope biology, including addressing questions of how fluidity is related to the disruption of lipid-related pathways and changes in drug accumulation. Larger-scale analyses could determine whether fluidity serves as a predictive marker of antibiotic susceptibility. Ultimately, this method provides a versatile framework for uncovering new insights into bacterial physiology and drug responses.

**Table 1.** Bacterial strains used in this study.

| Species | Strain | Genotype | Source | Reference |
| --- | --- | --- | --- | --- |
| *M. smegmatis* | mc<sup>2</sup>155 |  | ATCC 700084, gift of Dr. Eric Rubin |  |
| *M. smegmatis* | mc<sup>2</sup>155 | $\Delta cfa$ | Gift of Dr. Yasu Morita | Prithviraj et al., *mBio* 2023 |
| *M. smegmatis* | mc<sup>2</sup>155 | $\Delta lamA$ | Gift of Dr. Hesper Rego | Rego et al., *Nature* 2017 |
| *M. smegmatis* | mc<sup>2</sup>155 | $\Delta lprG-rv1410c$ | Gift of Dr. Eric Rubin | Farrow et al., *J Bacteriol.* 2007 |
| *M. smegmatis* | mc<sup>2</sup>155 | Pmop-<br>*luxABCDE-kanR* |  | Bai and Parkin et al., *ACS Infectious Diseases* 2020 |
| *M. tuberculosis* | H37Rv<br>mc<sup>2</sup>6020 | $\Delta panCD \Delta lysA$ | Gift of Dr. William Jacobs | |
| *M. abscessus* | 390R |  | Gift of Dr. Kyle Rhode |  |
| *M. abscessus* | 390S |  | Gift of Dr. Kyle Rhode |  |
| *M. avium*<br>subsp. *avium* | Chester |  | ATCC 25291; gift of Dr. Arthur Totten |  |
| *M. fortuitum* |  |  | ATCC 6841; gift of Dr. Arthur Totten |  |
| *M. scrofulaceum* |  |  | ATCC 19981; gift of Dr. Arthur Totten |  |
| *M. kansasii* |  |  | Gift of Dr. Arthur Totten |  |
| *Nocardia farcinica* | Trevisan |  | ATCC 3308; gift of Dr. Arthur Totten |  |

## METHODS

### Culture conditions

*Msm* cultures were incubated with shaking at 250 rpm; cultures for all other bacteria at 110 rpm. Unless otherwise indicated, all resuspension and wash steps for labeling experiments were performed with media and buffer prewarmed to 37 °C. For C-Laurdan labeling experiments in Figures 1 and 3-5, *Msm* strains were cultured in 7H9/Cas medium (Middlebrook 7H9 medium (HI MEDIA) with 1% w/v casamino acids (VWR), 0.2% w/v glucose, 0.05% v/v Tween 80). For Figure 2: *Mtb* was cultured in AMTB medium (Middlebrook 7H9 medium supplemented with 10% v/v oleic acid-albumin-dextrose-catalase (OADC) supplement (BD), 0.5% v/v glycerol, 0.2% w/v casamino acids (VWR J851), 80 mg/L L-lysine, 24 mg/L pantothenate, and 0.025% v/v Tyloxapol or on Middlebrook 7H10 agar (BD) supplemented with 10% v/v OADC, 0.5% v/v glycerol, 0.2% casamino acids, 80 mg/L L-lysine, 24 mg/L pantothenate). *Mycobacterium abscessus* (*Mab*) 390R, *Mab* 390S, *M. scrofulaceum, M. avium* subspecies *avium* Chester, *M. kansasii, M. fortuitum,* and *Msm* were cultured in 7H9/ADC (Middlebrook 7H9 medium supplemented with 10% albumin/dextrose/catalase (ADC), 0.5% glycerol, and 0.05% Tween 80). *N. farcinica* was cultured in 7H9/ADC and incubated at 30 °C. Unless otherwise noted, cells were labeled while in exponential phase (OD = 1-2, as determined in 7H9/Cas for *Msm*). To compare exponential and stationary phases, a single overnight starter culture was inoculated. Once the culture reached exponential phase (OD_₆₀₀_ = 1-2), a portion of the culture was diluted into fresh medium and grown for approximately one additional doubling before labeling. The remainder of the overnight culture was allowed to grow to stationary phase (OD_₆₀₀_ = 4-6), at which point it was harvested for labeling. For dose response curves in Figure 5, autoluminescent *Msm* was cultured in modified Sauton’s medium^81^ (4 g of L-asparagine, 0.5 g KH_2_PO_4_, 0.05 g ferric ammonium citrate, 0.5 g MgSO_4_•7H_2_O, 0.01 g ZnSO4, 2 g citric acid, 0.05% (v/v) Tyloxapol, pH 7 per 1L, plus a final concentration of 10 mM sodium propionate) and incubated at 37 °C. *Msm* was handled at BSL-1 containment; all other organisms were handled at BSL-2 containment.

### Labeling and preparation of bacteria for microscopy

Stocks of Laurdan (6-dodecanoyl-2-dimethylaminonaphthalene, Invitrogen), C-Laurdan (N-methyl-N-[6-(1-oxododecyl)-2-naphthalenyl]glycine, Setareh Biotech), and Pro12A^36^ [3-(N-dodecyl-2-(methyl(6-propionylnaphthalen-2-yl)amino)acetamido)propane-1-sulfonate, gift of Dr. Andrey Klymchenko] were prepared in DMSO. Exponential phase cultures (OD = 1-2) were concentrated to OD_600_= 5 by centrifugation at 9000 x*g* for 2.5 min and the decanted cell pellet was resuspended in growth medium plus membrane label to a final volume of 100 µL (final [DMSO] = 2-4%). Tubes were incubated in the dark at 37 °C at 250 rpm for 5 min, then washed once with PBS with 0.05% Tween 80 (PBST80). Cells were resuspended in 200 µL PBST80.

For imaging, agarose pads were prepared by dissolving 0.3 g agarose in 10 mL growth medium. For *Msm* 20 µL agarose solution was pipetted onto a glass slide, and covered with a coverslip. After ∼1-2 minutes, the coverslip was removed, 2 µL cell suspension pipetted onto the pad, the coverslip replaced, and the sandwich sealed with nail polish. For all other organisms, agarose pads were prepared by adding 90 µL to the well of a glass-bottom petri dish and covering with a coverslip. Petri dishes were transferred to the BSL-2 facility. Once labeled cells were ready for imaging, forceps were used to remove the coverslip and agar disk. 2 µL of labeled cells were added to the well of the glass-bottom dish, then forceps were used to place the agar disk and coverslip on top of the cells. Petri dishes were decontaminated with Vesphene II and sealed with parafilm prior to removal from BSL-2 containment and imaging.

### RADA labeling

Exponential phase *Msm* cells were concentrated to OD_600_ = 5 and resuspended in 7H9/Cas. RADA (Tocris Bioscience, 5 mM stock in DMSO) was added to a final concentration of 12.5 µM and incubated at 37 °C with shaking at 250 rpm for 15 min. After a 10-min incubation, C-Laurdan was added to a final concentration of 400 µM and cells were incubated for an additional 5 min. Cells were then washed and prepared for imaging as above.

### Confocal imaging

Images were acquired at 22 °C on a Zeiss LSM 980 Airyscan 2 NLO Two-Photon Confocal Microscope (Central Microscopy Imaging Center, Stony Brook University) with either a 100x phase contrast (Zeiss) or 100x differential interference contrast (Zeiss) objective. Samples were excited at 405 nm, and windows were set to simultaneously capture lipid-ordered and lipid-disordered emission peaks at 440 nm (420-460) and 490 nm (470-510), respectively. To allow comparison across samples within replicates, images were acquired using identical laser power and detector gain settings for each independent experiment.

### Generation and analysis of generalized polarization (GP) maps

Custom ImageJ plugins were generated to create whole cell, annular, and intracellular GP maps (code deposited in GitHub at https://github.com/jcslab-sbu). The plugins first concatenate all CZI images from a file folder into stacks according to their type (i.e., emission at 440 nm, at 490 nm, and white light image). Operations on the pixel intensities of the 440nm and 490 nm channels are then performed to create a GP map (ratio (I_440_-I_490_)/(I_440_+I_490_)) for every image in the folder. A threshold is then applied to the phase contrast stack (for phase contrast segmentation) or the 440 nm stack (for segmentation based on 440 nm emission): Background non-cellular signal is removed using a rolling-ball threshold, each image is converted to binary, and the nonzero pixel values are eroded. For the annular GP plugin, the edges of the cells are then identified and isolated and the rest of the stack is converted to a value of zero. For the intracellular GP plugin, the edges are eroded. Each image in the binary phase contrast or 440 nm stack is then multiplied by the GP map to isolate cell (or subcell)-associated GP values. To remove values contributed by background pixels, every pixel of each image is then iterated: If the pixel has a zero value, it is converted to NaN; if it is nonzero, it is divided by 255 (binary for the pixel intensity is 0-black or 255-white). The resulting stack of image files contains GP values only in areas of interest and invalidates background pixels so that the average across an area of interest is accurately reported. Finally, to exclude unlabeled cells from the phase contrast-segmented GP maps, the plugins overlay the stack of GP maps with a stack of 440 nm images. This step is unnecessary for the 440 nm-segmented GP maps, as unlabeled cells are excluded during the segmentation step.

After annular, whole cell, and intracellular GP maps were generated, average GP was measured for individual cells and regions of interest using the freehand selection tool and the measure function. For high vs. low bilateral GP analysis in Figure 1, annular GP maps were generated, cells were manually bisected along the long axis, and the freehand selection tool was used to measure the GP at each side. The paired values were then sorted by high-GP vs. low-GP sides for each cell. For polar and sidewall analysis of annular GP, polar GP was averaged across ∼15% of the cell at each end, and sidewall GP was measured in the remaining central ∼70%, similar to a previous report^38^. For RADA experiments, the GP maps were overlaid with the RADA signal. For RADA-bright vs. RADA-dim polar analysis, annular GP at the poles was averaged as before across ∼15% of cell length at each end and indexed as at the RADA-bright or RADA-dim pole. For septal analysis, the overlaid RADA signal was used to guide the collection of GP data at the septa and cell periphery from whole cell GP maps using a 3-pixel wide freehand line. For demographs and line profiles, whole cell GP maps were generated and line profiles were collected from labeled cells by drawing a 40-pixel wide straight line through the cell along the long axis. Line profiles were saved as CSV files into folders for each condition.

Demographs were generated using a custom MATLAB script that imports folders of CSV files containing line profiles, removes non-numerical (NaN) values, and sorts each line profile by length. Profiles are aligned by the high-GP pole and centered at the midpoint. Demographs are generated by stacking aligned profiles, with the x-axis representing distance from the cell midpoint (µm) and the y-axis corresponding to individual cells ordered by length. Intensity values for GP were normalized and scaled to a fixed color axis, with zeros rendered transparent. Average line profiles were generated by using a custom MATLAB script to align the cells by the high-GP pole and normalize each profile to percent cell length (0–100%), interpolate onto a shared axis using linear interpolation, and smooth the resulting profile using the Lowess method. The solid line represents the mean profile across all cells and the shaded region indicates ±1 standard deviation.

### Flow cytometry

*Msm* was labeled with 400 µM C-Laurdan as above in “Labeling and preparation of bacteria for microscopy”, but washed with filtered PBS (instead of PBST80) and resuspended in 1 mL filtered PBS. Flow cytometry was performed on a Beckman Coulter CytoFLEX Flow Cytometer (Stony Brook University Flow Cytometry Research Core Facility). Cells were excited at 375 nm and emission intensity collected for 20,000 events using bandpass filters at 450/45 nm (428-473 nm) and 525/40 nm (505-545 nm). Events were gated for live, C-Laurdan-positive cells using unlabeled controls, and GP was calculated using the modified formula GP=(I_450_ – I_525_)/(I_450_ – I_525_).

### BSA extraction of C-Laurdan

*Msm* was labeled with 400 µM C-Laurdan as above in “Labeling and preparation of bacteria for microscopy”. Cells were pelleted by centrifugation at 9000 x*g* for 2.5 min. Decanted pellets were resuspended in 200 µL PBS or PBS with 5 mg/mL fatty acid free bovine serum albumin (BSA) and incubated for 20 min at 37 °C, 250 rpm. Cells were then pelleted and resuspended in 1 mL of filtered PBS for analysis as above in “Flow cytometry”.

### Antibiotic treatment for GP analysis

Exponential phase *Msm* was diluted to OD_600_ = ∼0.5 (WT) or OD_600_ = ∼0.7 (Δ*lprG-rv1410c;* higher OD than WT to account for a mild growth defect) in 7H9/Cas with various antibiotics: 2 and 50 µM ethambutol (1x and 20x MIC), 1 and 20 µM hygromycin (1x and 20x MIC), or 5 mM (20x MIC) vancomycin. Antibiotic stocks were dissolved in autoclaved ultrapure water, which served as the vehicle control. The samples were incubated at 37 °C, 250 rpm for 3 hrs. The OD_600_ was measured and cells were labeled with 400 µM C-Laurdan as above in “Labeling and preparation of bacteria for microscopy”.

### Dose response curves

Assays were performed as previously reported. Briefly, overnight cultures of autoluminescent *Msm*^81^ in modified Sauton’s medium were grown to OD_600_ ∼0.8-1, then subcultured to OD_600_ 0.1 and incubated with varying concentrations of vancomycin, hygromycin, and/or ethambutol in a total volume of 200 μL in white clear bottom 96-well plates (Corning). Plates were incubated in a multimode plate reader (FilterMax F5, Molecular Devices) at 37 °C with 15 min orbital shaking between measurements for 19 hours total (4-5 doubling times in modified Sauton’s medium). Percent viability was calculated by normalizing RLUs from antibiotic-treated cells to untreated controls: RLU(+cmpd)/RLU(−cmpd) x 100. Dose-response curves were fit in GraphPad Prism using nonlinear regression with a four-parameter logistic equation: Y = Bottom + (Top – Bottom) / (1 + 10^((LogIC50 – X) * HillSlope)). For normalized data, Top and Bottom were constrained to 100 and 0, respectively.

## ACKNOWLEDGEMENTS

We thank Julian Maceren for contributions to preliminary experiments and the Seeliger lab for helpful discussions. We are grateful to Hesper Rego, Henrik Strahl, and Erwin London for valuable input, and to Hesper Rego, Yasu Morita, Andrey Klymchenko, Kyle Rhode, and Arthur Totten for providing strains and reagents. This work was supported by F30 AI179077 (I.S.), R35 GM119437 (J.C.S.), T32 GM008444 (I.S.), T32 AI007539 (I.S. and L.P.), and T32 GM136572 (L.P.).

## REFERENCES

(1) Rossignol, M.; Thomas, P.; Grignon, C. Proton Permeability of Liposomes from Natural Phospholipid Mixtures. Biochim. Biophys. Acta BBA - Biomembr. 1982, 684 (2), 195–199. 10.1016/0005-2736(82)90005-0.

(2) van de Vossenberg, J. L. C. M.; Driessen, A. J. M.; da Costa, M. S.; Konings, W. N. Homeostasis of the Membrane Proton Permeability in *Bacillus Subtilis* Grown at Different Temperatures. Biochim. Biophys. Acta BBA - Biomembr. 1999, 1419 (1), 97–104. 10.1016/S0005-2736(99)00063-2.

(3) Lenaz, G. Lipid Fluidity and Membrane Protein Dynamics. Biosci. Rep. 1987, 7 (11), 823–837. 10.1007/BF01119473.

(4) Pande, A. H.; Qin, S.; Tatulian, S. A. Membrane Fluidity Is a Key Modulator of Membrane Binding, Insertion, and Activity of 5-Lipoxygenase. Biophys. J. 2005, 88 (6), 4084–4094. 10.1529/biophysj.104.056788.

(5) Budin, I.; de Rond, T.; Chen, Y.; Chan, L. J. G.; Petzold, C. J.; Keasling, J. D. Viscous Control of Cellular Respiration by Membrane Lipid Composition. Science 2018, 362 (6419), 1186–1189. 10.1126/science.aat7925.

(6) Gohrbandt, M.; Lipski, A.; Grimshaw, J. W.; Buttress, J. A.; Baig, Z.; Herkenhoff, B.; Walter, S.; Kurre, R.; Deckers-Hebestreit, G.; Strahl, H. Low Membrane Fluidity Triggers Lipid Phase Separation and Protein Segregation in Living Bacteria. EMBO J. 2022, 41 (5), e109800. 10.15252/embj.2021109800.

(7) Podoll, J. D.; Rosen, E.; Wang, W.; Gao, Y.; Zhang, J.; Wang, X. A Small-Molecule Membrane Fluidizer Re-Sensitizes Methicillin-Resistant Staphylococcus Aureus (MRSA) to β-Lactam Antibiotics. Antimicrob. Agents Chemother. 2023, 67 (10), e00051–23. 10.1128/aac.00051-23.

(8) Bessa, L. J.; Ferreira, M.; Gameiro, P. Evaluation of Membrane Fluidity of Multidrug-Resistant Isolates of *Escherichia Coli* and *Staphylococcus Aureus* in Presence and Absence of Antibiotics. J. Photochem. Photobiol. B 2018, 181, 150–156. 10.1016/j.jphotobiol.2018.03.002.

(9) Müller, A.; Wenzel, M.; Strahl, H.; Grein, F.; Saaki, T. N. V.; Kohl, B.; Siersma, T.; Bandow, J. E.; Sahl, H.-G.; Schneider, T.; Hamoen, L. W. Daptomycin Inhibits Cell Envelope Synthesis by Interfering with Fluid Membrane Microdomains. Proc. Natl. Acad. Sci. U. S. A. 2016, 113 (45), E7077–E7086. 10.1073/pnas.1611173113.

(10) Saeloh, D.; Tipmanee, V.; Jim, K. K.; Dekker, M. P.; Bitter, W.; Voravuthikunchai, S. P.; Wenzel, M.; Hamoen, L. W. The Novel Antibiotic Rhodomyrtone Traps Membrane Proteins in Vesicles with Increased Fluidity. PLoS Pathog. 2018, 14 (2), e1006876. 10.1371/journal.ppat.1006876.

(11) Nikaido, H.; Kim, S.-H.; Rosenberg, E. Y. Physical Organization of Lipids in the Cell Wall of Mycobacterium Chelonae. Mol. Microbiol. 1993, 8 (6), 1025–1030. 10.1111/j.1365-2958.1993.tb01647.x.

(12) Liu, J.; Barry, C. E.; Besra, G. S.; Nikaido, H. Mycolic Acid Structure Determines the Fluidity of the Mycobacterial Cell Wall*. J. Biol. Chem. 1996, 271 (47), 29545–29551. 10.1074/jbc.271.47.29545.

(13) Liu, J.; Rosenberg, E. Y.; Nikaido, H. Fluidity of the Lipid Domain of Cell Wall from Mycobacterium Chelonae. Proc. Natl. Acad. Sci. 1995, 92 (24), 11254–11258. 10.1073/pnas.92.24.11254.

(14) Liu, J.; Nikaido, H. A Mutant of Mycobacterium Smegmatis Defective in the Biosynthesis of Mycolic Acids Accumulates Meromycolates. Proc. Natl. Acad. Sci. 1999, 96 (7), 4011–4016. 10.1073/pnas.96.7.4011.

(15) Schami, A.; Islam, M. N.; Belisle, J. T.; Torrelles, J. B. Drug-Resistant Strains of Mycobacterium Tuberculosis: Cell Envelope Profiles and Interactions with the Host. Front. Cell. Infect. Microbiol. 2023, 13. 10.3389/fcimb.2023.1274175.

(16) Xu, W.; DeJesus, M. A.; Rücker, N.; Engelhart, C. A.; Wright, M. G.; Healy, C.; Lin, K.; Wang, R.; Park, S. W.; Ioerger, T. R.; Schnappinger, D.; Ehrt, S. Chemical Genetic Interaction Profiling Reveals Determinants of Intrinsic Antibiotic Resistance in Mycobacterium Tuberculosis. Antimicrob. Agents Chemother. 2017, 61 (12), e01334–17. 10.1128/AAC.01334-17.

(17) Bellerose, M. M.; Proulx, M. K.; Smith, C. M.; Baker, R. E.; Ioerger, T. R.; Sassetti, C. M. Distinct Bacterial Pathways Influence the Efficacy of Antibiotics against *Mycobacterium Tuberculosis*. mSystems 2020, 5 (4), e00396–20, /msystems/5/4/msys.00396-20.atom. 10.1128/mSystems.00396-20.

(18) Li, S.; Poulton, N. C.; Chang, J. S.; Azadian, Z. A.; DeJesus, M. A.; Ruecker, N.; Zimmerman, M. D.; Eckartt, K. A.; Bosch, B.; Engelhart, C. A.; Sullivan, D. F.; Gengenbacher, M.; Dartois, V. A.; Schnappinger, D.; Rock, J. M. CRISPRi Chemical Genetics and Comparative Genomics Identify Genes Mediating Drug Potency in Mycobacterium Tuberculosis. Nat. Microbiol. 2022, 7 (6), 766–779. 10.1038/s41564-022-01130-y.

(19) Koh, E.-I.; Oluoch, P. O.; Ruecker, N.; Proulx, M. K.; Soni, V.; Murphy, K. C.; Papavinasasundaram, K.; Reames, C. J.; Trujillo, C.; Zaveri, A.; Zimmerman, M. D.; Aslebagh, R.; Baker, R. E.; Shaffer, S. A.; Guinn, K. M.; Fitzgerald, M.; Dartois, V.; Ehrt, S.; Hung, D. T.; Ioerger, T. R.; Rubin, E. J.; Rhee, K. Y.; Schnappinger, D.; Sassetti, C. M. Chemical–Genetic Interaction Mapping Links Carbon Metabolism and Cell Wall Structure to Tuberculosis Drug Efficacy. Proc. Natl. Acad. Sci. 2022, 119 (15), e2201632119. 10.1073/pnas.2201632119.

(20) Poulton, N. C.; Rock, J. M. Unraveling the Mechanisms of Intrinsic Drug Resistance in Mycobacterium Tuberculosis. Front. Cell. Infect. Microbiol. 2022, 12. 10.3389/fcimb.2022.997283.

(21) Adhyapak, P.; Srivatsav, A. T.; Mishra, M.; Singh, A.; Narayan, R.; Kapoor, S. Dynamical Organization of Compositionally Distinct Inner and Outer Membrane Lipids of Mycobacteria. Biophys. J. 2020, 118 (6), 1279–1291. 10.1016/j.bpj.2020.01.027.

(22) Brown, C. M.; Corey, R. A.; Grélard, A.; Gao, Y.; Choi, Y. K.; Luna, E.; Gilleron, M.; Destainville, N.; Nigou, J.; Loquet, A.; Fullam, E.; Im, W.; Stansfeld, P. J.; Chavent, M. Supramolecular Organization and Dynamics of Mannosylated Phosphatidylinositol Lipids in the Mycobacterial Plasma Membrane. Proc. Natl. Acad. Sci. 2023, 120 (5), e2212755120. 10.1073/pnas.2212755120.

(23) Parasassi, T.; Krasnowska, E. K.; Bagatolli, L.; Gratton, E. Laurdan and Prodan as Polarity-Sensitive Fluorescent Membrane Probes. J. Fluoresc. 1998, 8 (4), 365–373. 10.1023/A:1020528716621.

(24) Klymchenko, A. S. Fluorescent Probes for Lipid Membranes: From the Cell Surface to Organelles. Acc. Chem. Res. 2023, 56 (1), 1–12. 10.1021/acs.accounts.2c00586.

(25) Qin, X.; Yang, X.; Du, L.; Li, M. Polarity-Based Fluorescence Probes: Properties and Applications. RSC Med. Chem. 12 (11), 1826–1838. 10.1039/d1md00170a.

(26) Klymchenko, A. S. Solvatochromic and Fluorogenic Dyes as Environment-Sensitive Probes: Design and Biological Applications. Acc. Chem. Res. 2017, 50 (2), 366–375. 10.1021/acs.accounts.6b00517.

(27) Christensen, H.; Garton, N. J.; Horobin, R. W.; Minnikin, D. E.; Barer, M. R. Lipid Domains of Mycobacteria Studied with Fluorescent Molecular Probes. Mol. Microbiol. 1999, 31 (5), 1561–1572. 10.1046/j.1365-2958.1999.01304.x.

(28) Kamariza, M.; Keyser, S. G. L.; Utz, A.; Knapp, B. D.; Ealand, C.; Ahn, G.; Cambier, C. J.; Chen, T.; Kana, B.; Huang, K. C.; Bertozzi, C. R. Toward Point-of-Care Detection of Mycobacterium Tuberculosis: A Brighter Solvatochromic Probe Detects Mycobacteria within Minutes. JACS Au 2021, 1 (9), 1368–1379. 10.1021/jacsau.1c00173.

(29) Kamariza, M.; Shieh, P.; Bertozzi, C. R. Imaging Mycobacterial Trehalose Glycolipids. In Methods in Enzymology; Elsevier, 2018; Vol. 598, pp 355–369. 10.1016/bs.mie.2017.09.002.

(30) Gunther, G.; Malacrida, L.; Jameson, D. M.; Gratton, E.; Sánchez, S. A. LAURDAN since Weber: The Quest for Visualizing Membrane Heterogeneity. Acc. Chem. Res. 2021, 54 (4), 976–987. 10.1021/acs.accounts.0c00687.

(31) Sullivan, M. R.; McGowen, K.; Liu, Q.; Akusobi, C.; Young, D. C.; Mayfield, J. A.; Raman, S.; Wolf, I. D.; Moody, D. B.; Aldrich, C. C.; Muir, A.; Rubin, E. J. Biotin-Dependent Cell Envelope Remodeling Is Required for Mycobacterium Abscessus Survival in Lung Infection. Nat. Microbiol. 2023, 8 (3), 481–497. 10.1038/s41564-022-01307-5.

(32) Aldridge, B. B.; Fernandez-Suarez, M.; Heller, D.; Ambravaneswaran, V.; Irimia, D.; Toner, M.; Fortune, S. Asymmetry and Aging of Mycobacterial Cells Leads to Variable Growth and Antibiotic Susceptibility. Science 2012, 335 (6064), 100–104. 10.1126/science.1216166.

(33) Rego, E. H.; Audette, R. E.; Rubin, E. J. Deletion of a Mycobacterial Divisome Factor Collapses Single-Cell Phenotypic Heterogeneity. Nature 2017, 546 (7656), 153–157. 10.1038/nature22361.

(34) Kim, H. M.; Choo, H.; Jung, S.; Ko, Y.; Park, W.; Jeon, S.; Kim, C. H.; Joo, T.; Cho, B. R. A Two-Photon Fluorescent Probe for Lipid Raft Imaging: C-Laurdan. ChemBioChem 2007, 8 (5), 553–559. 10.1002/cbic.200700003.

(35) Barucha-Kraszewska, J.; Kraszewski, S.; Ramseyer, C. Will C-Laurdan Dethrone Laurdan in Fluorescent Solvent Relaxation Techniques for Lipid Membrane Studies? Langmuir 2013, 29 (4), 1174–1182. 10.1021/la304235r.

(36) Danylchuk, D. I.; Sezgin, E.; Chabert, P.; Klymchenko, A. S. Redesigning Solvatochromic Probe Laurdan for Imaging Lipid Order Selectively in Cell Plasma Membranes. Anal. Chem. 2020, 92 (21), 14798–14805. 10.1021/acs.analchem.0c03559.

(37) Zhu, J.; Wolf, I. D.; Dulberger, C. L.; Won, H. I.; Kester, J. C.; Judd, J. A.; Wirth, S. E.; Clark, R. R.; Li, Y.; Luo, Y.; Gray, T. A.; Wade, J. T.; Derbyshire, K. M.; Fortune, S. M.; Rubin, E. J. Spatiotemporal Localization of Proteins in Mycobacteria. Cell Rep. 2021, 37 (13), 110154. 10.1016/j.celrep.2021.110154.

(38) García-Heredia, A.; Pohane, A. A.; Melzer, E. S.; Carr, C. R.; Fiolek, T. J.; Rundell, S. R.; Lim, H. C.; Wagner, J. C.; Morita, Y. S.; Swarts, B. M.; Siegrist, M. S. Peptidoglycan Precursor Synthesis along the Sidewall of Pole-Growing Mycobacteria. eLife 2018, 7, e37243. 10.7554/eLife.37243.

(39) Neidhardt, F. C. Escherichia Coli and Salmonella: Cellular and Molecular Biology; ASM Press, 1996.

(40) Smeulders, M. J.; Keer, J.; Speight, R. A.; Williams, H. D. Adaptation of Mycobacterium Smegmatis to Stationary Phase. J. Bacteriol. 1999, 181 (1), 270–283. 10.1128/jb.181.1.270-283.1999.

(41) Sartain, M. J.; Dick, D. L.; Rithner, C. D.; Crick, D. C.; Belisle, J. T. Lipidomic Analyses of Mycobacterium Tuberculosis Based on Accurate Mass Measurements and the Novel “Mtb LipidDB.” J. Lipid Res. 2011, 52 (5), 861. 10.1194/jlr.M010363.

(42) Laydevant, F.; Mahabadi, M.; Llido, P.; Bourgouin, J.-P.; Caron, L.; Arnold, A. A.; Marcotte, I.; Warschawski, D. E. Growth-Phase Dependence of Bacterial Membrane Lipid Profile and Labeling for in-Cell Solid-State NMR Applications. Biochim. Biophys. Acta BBA - Biomembr. 2022, 1864 (2), 183819. 10.1016/j.bbamem.2021.183819.

(43) P. Menon, A.; Lee, T.-H.; Aguilar, M.-I.; Kapoor, S. Decoding the Role of Mycobacterial Lipid Remodelling and Membrane Dynamics in Antibiotic Tolerance. Chem. Sci. 2024, 15 (45), 19084–19093. 10.1039/D4SC06618A.

(44) Hobby, G. L.; Lenert, T. F. The in Vitro Action of Antituberculous Agents Against Multiplying and Nonmultiplying Microbial Cells,. Am. Rev. Tuberc. Pulm. Dis. 1957, 76 (6), 1031–1048. 10.1164/artpd.1957.76.6.1031.

(45) Hammond, R. J. H.; Baron, V. O.; Oravcova, K.; Lipworth, S.; Gillespie, S. H. Phenotypic Resistance in Mycobacteria: Is It Because I Am Old or Fat That I Resist You? J. Antimicrob. Chemother. 2015, 70 (10), 2823–2827. 10.1093/jac/dkv178.

(46) Gomez, J. E.; McKinney, J. D. M. Tuberculosis Persistence, Latency, and Drug Tolerance. Tuberculosis 2004, 84 (1–2), 29–44. 10.1016/j.tube.2003.08.003.

(47) Thanky, N. R.; Young, D. B.; Robertson, B. D. Unusual Features of the Cell Cycle in Mycobacteria: Polar-Restricted Growth and the Snapping-Model of Cell Division. Tuberculosis 2007, 87 (3), 231–236. 10.1016/j.tube.2006.10.004.

(48) Bach, J. N.; Bramkamp, M. Flotillins Functionally Organize the Bacterial Membrane. Mol. Microbiol. 2013, 88 (6), 1205–1217. 10.1111/mmi.12252.

(49) Strahl, H.; Bürmann, F.; Hamoen, L. W. The Actin Homologue MreB Organizes the Bacterial Cell Membrane. Nat. Commun. 2014, 5 (1), 3442. 10.1038/ncomms4442.

(50) Prithviraj, M.; Kado, T.; Mayfield, J. A.; Young, D. C.; Huang, A. D.; Motooka, D.; Nakamura, S.; Siegrist, M. S.; Moody, D. B.; Morita, Y. S. Tuberculostearic Acid Controls Mycobacterial Membrane Compartmentalization. mBio 14 (2), e03396–22. 10.1128/mbio.03396-22.

(51) Meniche, X.; Otten, R.; Siegrist, M. S.; Baer, C. E.; Murphy, K. C.; Bertozzi, C. R.; Sassetti, C. M. Subpolar Addition of New Cell Wall Is Directed by DivIVA in Mycobacteria. Proc. Natl. Acad. Sci. U. S. A. 2014, 111 (31), E3243–E3251. 10.1073/pnas.1402158111.

(52) Meyer, F. M.; Bramkamp, M. Cell Wall Synthesizing Complexes in *Mycobacteriales*. Curr. Opin. Microbiol. 2024, 79, 102478. 10.1016/j.mib.2024.102478.

(53) Morita, Y. S.; Velasquez, R.; Taig, E.; Waller, R. F.; Patterson, J. H.; Tull, D.; Williams, S. J.; Billman-Jacobe, H.; McConville, M. J. Compartmentalization of Lipid Biosynthesis in Mycobacteria. J. Biol. Chem. 2005, 280 (22), 21645–21652. 10.1074/jbc.M414181200.

(54) Hayashi, J. M.; Luo, C.-Y.; Mayfield, J. A.; Hsu, T.; Fukuda, T.; Walfield, A. L.; Giffen, S. R.; Leszyk, J. D.; Baer, C. E.; Bennion, O. T.; Madduri, A.; Shaffer, S. A.; Aldridge, B. B.; Sassetti, C. M.; Sandler, S. J.; Kinoshita, T.; Moody, D. B.; Morita, Y. S. Spatially Distinct and Metabolically Active Membrane Domain in Mycobacteria. Proc. Natl. Acad. Sci. U. S. A. 2016, 113 (19), 5400–5405. 10.1073/pnas.1525165113.

(55) Meniche, X.; Otten, R.; Siegrist, M. S.; Baer, C. E.; Murphy, K. C.; Bertozzi, C. R.; Sassetti, C. M. Subpolar Addition of New Cell Wall Is Directed by DivIVA in Mycobacteria. Proc. Natl. Acad. Sci. U. S. A. 2014, 111 (31), E3243–E3251. 10.1073/pnas.1402158111.

(56) Liu, D.; Hao, K.; Wang, W.; Peng, C.; Dai, Y.; Jin, R.; Xu, W.; He, L.; Wang, H.; Wang, H.; Zhang, L.; Wang, Q. Rv2629 Overexpression Delays Mycobacterium Smegmatis and Mycobacteria Tuberculosis Entry into Log-Phase and Increases Pathogenicity of Mycobacterium Smegmatis in Mice. Front. Microbiol. 2017, *8*, 2231. 10.3389/fmicb.2017.02231.

(57) Piddock, L. J. V.; Williams, K. J.; Ricci, V. Accumulation of Rifampicin by Mycobacterium Aurum, Mycobacterium Smegmatis and Mycobacterium Tuberculosis. J. Antimicrob. Chemother. 2000, 45 (2), 159–165. 10.1093/jac/45.2.159.

(58) Flores, A. R.; Parsons, L. M.; Pavelka, M. S. Characterization of Novel Mycobacterium Tuberculosis and Mycobacterium Smegmatis Mutants Hypersusceptible to β-Lactam Antibiotics. J. Bacteriol. 2005, 187 (6), 1892–1900. 10.1128/JB.187.6.1892-1900.2005.

(59) Bigi, F.; Gioffré, A.; Klepp, L.; de la Paz Santangelo, M.; Alito, A.; Caimi, K.; Meikle, V.; Zumárraga, M.; Taboga, O.; Romano, M. I.; Cataldi, A. The Knockout of the *lprG-Rv1410* Operon Produces Strong Attenuation of *Mycobacterium Tuberculosis*. Microbes Infect. 2004, 6 (2), 182–187. 10.1016/j.micinf.2003.10.010.

(60) Martinot, A. J.; Farrow, M.; Bai, L.; Layre, E.; Cheng, T.-Y.; Tsai, J. H.; Iqbal, J.; Annand, J. W.; Sullivan, Z. A.; Hussain, M. M.; Sacchettini, J.; Moody, D. B.; Seeliger, J. C.; Rubin, E. J. Mycobacterial Metabolic Syndrome: LprG and Rv1410 Regulate Triacylglyceride Levels, Growth Rate and Virulence in Mycobacterium Tuberculosis. PLOS Pathog. 2016, 12 (1), e1005351. 10.1371/journal.ppat.1005351.

(61) Bianco, M. V.; Blanco, F. C.; Imperiale, B.; Forrellad, M. A.; Rocha, R. V.; Klepp, L. I.; Cataldi, A. A.; Morcillo, N.; Bigi, F. Role of P27-P55 Operon from Mycobacterium Tuberculosis in the Resistance to Toxic Compounds. BMC Infect. Dis. 2011, 11 (1), 195. 10.1186/1471-2334-11-195.

(62) Farrow, M. F.; Rubin, E. J. Function of a Mycobacterial Major Facilitator Superfamily Pump Requires a Membrane-Associated Lipoprotein. J. Bacteriol. 2008, 190 (5), 1783–1791. 10.1128/JB.01046-07.

(63) Hohl, M.; Remm, S.; Eskandarian, H. A.; Dal Molin, M.; Arnold, F. M.; Hürlimann, L. M.; Krügel, A.; Fantner, G. E.; Sander, P.; Seeger, M. A. Increased Drug Permeability of a Stiffened Mycobacterial Outer Membrane in Cells Lacking MFS Transporter Rv1410 and Lipoprotein LprG. Mol. Microbiol. 2019, 111 (5), 1263–1282. 10.1111/mmi.14220.

(64) Ayala-Torres, C.; Hernández, N.; Galeano, A.; Novoa-Aponte, L.; Soto, C.-Y. Zeta Potential as a Measure of the Surface Charge of Mycobacterial Cells. Ann. Microbiol. 2014, 64 (3), 1189–1195. 10.1007/s13213-013-0758-y.

(65) Wong, A. M.; Budin, I. Organelle-Targeted Laurdans Measure Heterogeneity in Subcellular Membranes and Their Responses to Saturated Lipid Stress. ACS Chem. Biol. 2024, 19 (8), 1773–1785. 10.1021/acschembio.4c00249.

(66) Mallick, I.; Santucci, P.; Poncin, I.; Point, V.; Kremer, L.; Cavalier, J.-F.; Canaan, S. Intrabacterial Lipid Inclusions in Mycobacteria: Unexpected Key Players in Survival and Pathogenesis? FEMS Microbiol. Rev. 2021, 45 (6), fuab029. 10.1093/femsre/fuab029.

(67) Bansal-Mutalik, R.; Nikaido, H. Quantitative Lipid Composition of Cell Envelopes of Corynebacterium Glutamicum Elucidated through Reverse Micelle Extraction. Proc. Natl. Acad. Sci. 2011, 108 (37), 15360–15365. 10.1073/pnas.1112572108.

(68) Los, D. A.; Murata, N. Membrane Fluidity and Its Roles in the Perception of Environmental Signals. Biochim. Biophys. Acta BBA - Biomembr. 2004, 1666 (1), 142–157. 10.1016/j.bbamem.2004.08.002.

(69) Sezgin, E.; Levental, I.; Mayor, S.; Eggeling, C. The Mystery of Membrane Organization: Composition, Regulation and Roles of Lipid Rafts. Nat. Rev. Mol. Cell Biol. 2017, 18 (6), 361–374. 10.1038/nrm.2017.16.

(70) Bachmann, N. L.; Salamzade, R.; Manson, A. L.; Whittington, R.; Sintchenko, V.; Earl, A. M.; Marais, B. J. Key Transitions in the Evolution of Rapid and Slow Growing Mycobacteria Identified by Comparative Genomics. Front. Microbiol. 2020, 10, 3019. 10.3389/fmicb.2019.03019.

(71) Minnikin, D. E.; Minnikin, S. M.; Parlett, J. H.; Goodfellow, M.; Magnusson, M. Mycolic Acid Patterns of Some Species of Mycobacterium. Arch. Microbiol. 1984, 139–139 (2–3), 225–231. 10.1007/BF00402005.

(72) Rodriguez-Rivera, F. P.; Zhou, X.; Theriot, J. A.; Bertozzi, C. R. Visualization of Mycobacterial Membrane Dynamics in Live Cells. J. Am. Chem. Soc. 2017, 139 (9), 3488– 3495. 10.1021/jacs.6b12541.

(73) Daffé, M.; Draper, P. The Envelope Layers of Mycobacteria with Reference to Their Pathogenicity. In Advances in Microbial Physiology; Poole, R. K., Ed.; Academic Press, 1997; Vol. 39, pp 131–203. 10.1016/S0065-2911(08)60016-8.

(74) Chiaradia, L.; Lefebvre, C.; Parra, J.; Marcoux, J.; Burlet-Schiltz, O.; Etienne, G.; Tropis, M.; Daffé, M. Dissecting the Mycobacterial Cell Envelope and Defining the Composition of the Native Mycomembrane. Sci. Rep. 2017, 7 (1), 12807. 10.1038/s41598-017-12718-4.

(75) Rahlwes, K. C.; Sparks, I. L.; Morita, Y. S. Cell Walls and Membranes of Actinobacteria. In Bacterial Cell Walls and Membranes; Kuhn, A., Ed.; Springer International Publishing: Cham, 2019; Vol. 92, pp 417–469. 10.1007/978-3-030-18768-2_13.

(76) Wuo, M. G.; Dulberger, C. L.; Warner, T. C.; Brown, R. A.; Sturm, A.; Ultee, E.; Bloom-Ackermann, Z.; Choi, C.; Zhu, J.; Garner, E. C.; Briegel, A.; Hung, D. T.; Rubin, E. J.; Kiessling, L. L. Fluorogenic Probes of the Mycobacterial Membrane as Reporters of Antibiotic Action. J. Am. Chem. Soc. 2024, 146 (26), 17669–17678. 10.1021/jacs.4c00617.

(77) Smelyansky, S. R.; Ma, C.-W.; Marando, V. M.; Babunovic, G. H.; Lee, S. Y.; Bryson, B. D.; Kiessling, L. L. Exploiting Thioether Reactivity to Label Mycobacterial Glycans. Proc. Natl. Acad. Sci. 2025, 122 (19), e2422185122. 10.1073/pnas.2422185122.

(78) Zhou, X.; Rodriguez-Rivera, F. P.; Lim, H. C.; Bell, J. C.; Bernhardt, T. G.; Bertozzi, C. R.; Theriot, J. A. Sequential Assembly of the Septal Cell Envelope Prior to V Snapping in Corynebacterium Glutamicum. Nat. Chem. Biol. 2019, 15 (3), 221–231. 10.1038/s41589-018-0206-1.

(79) Icha, J.; Weber, M.; Waters, J. C.; Norden, C. Phototoxicity in Live Fluorescence Microscopy, and How to Avoid It. BioEssays 2017, 39 (8), 1700003. 10.1002/bies.201700003.

(80) Efremov, Y. M.; Shimolina, L.; Gulin, A.; Ignatova, N.; Gubina, M.; Kuimova, M. K.; Timashev, P. S.; Shirmanova, M. V. Correlation of Plasma Membrane Microviscosity and Cell Stiffness Revealed via Fluorescence-Lifetime Imaging and Atomic Force Microscopy. Cells 2023, 12 (21), 2583. 10.3390/cells12212583.

(81) Bai, L.; Parkin, L. A.; Zhang, H.; Shum, R.; Previti, M. L.; Seeliger, J. C. Dimethylaminophenyl Hydrazides as Inhibitors of the Lipid Transport Protein LprG in Mycobacteria. ACS Infect. Dis. 2020, 6 (4), 637–648. 10.1021/acsinfecdis.9b00497.

